# An updated and expanded characterization of the biological sciences academic job market

**DOI:** 10.1101/2024.07.31.606033

**Authors:** Brooklynn Flynn, Ariangela J. Kozik, You Cheng, Ada K. Hagan, Jennifer Ng, Christopher T. Smith, Amanda Haage, Nafisa M. Jadavji

## Abstract

In the biological sciences, many areas of uncertainty exist regarding the factors that contribute to success within the faculty job market. Earlier work from our group reported that beyond certain thresholds, academic and career metrics like the number of publications, fellowships or career transition awards, and years of experience did not separate applicants who received job offers from those who did not. Questions still exist regarding how academic and professional achievements influence job offers and if candidate demographics differentially influence outcomes. To continue addressing these gaps, we initiated surveys collecting data from faculty applicants in the biological sciences field for three hiring cycles in North America (Fall 2019 to the end of May 2022), a total of 449 respondents were included in our analysis. These responses highlight the interplay between various scholarly metrics, extensive demographic information, and hiring outcomes, and for the first time, allowed us to look at persons historically excluded due to ethnicity or race (PEER) status in the context of the faculty job market. Between 2019 and 2022, we found that the number of applications submitted, position seniority, and identifying as a women or transgender were positively correlated with a faculty job offer. Applicant age, residence, first generation status, and number of postdocs, however, were negatively correlated with receiving a faculty job offer. Our data are consistent with other surveys that also highlight the influence of achievements and other factors in hiring processes. Providing baseline comparative data for job seekers can support their informed decision-making in the market and is a first step towards demystifying the faculty job market.

## 1 Introduction

Landing a faculty position in the biomedical fields is competitive and full of challenges due to the job market dynamics, personal characteristics, and the negative effects of the pandemic (1–3). In biomedical fields, the number of PhD holders increases each year, but faculty positions have remained stagnant (4). Estimates suggest that there is only one tenure track faculty position available for every 6.3 PhD graduates (5). These dynamics have led to increased pressure on prospective faculty entering the job market and shifting sentiments among PhD holders in deciding if this is a career they want to pursue (6). To compete for faculty positions, PhD holders are increasingly compelled to stay in low paying ‘training’ positions (i.e. postdoctoral positions) for extended periods (7,8). Additionally, there has been a decrease in the number of PhD holders with an interest in completing a postdoc altogether, due to specific job attributes, economic stressors, and the PhD holder’s perceptions of their own research (9). Our research on the faculty job market is interested in determining factors that lead to success on the faculty job market given this climate (2,10,11).

Since conducting our first survey in 2018 (11), we have aimed to expand our research to investigate diversity in the faculty job market. The diversity of biomedical sciences PhD holders is not reflected in the make-up of current faculty members and the number of underrepresented minority (URM) candidates hired each year has decreased (12). Furthermore, there have been studies demonstrating that ethnicity and race affect the job offers of PhDs (13–15). To expand our analysis of factors that lead to success on the biological sciences faculty job market, in the present study we specifically asked PhD holder whether they were classified as persons excluded because of their ethnicity or race (PEER) (13). Including PEER status will allow our research to start characterizing this population of PhD holders, since historic data has shown that applicants who were white, male, and gender-conforming received a disproportionate number of faculty job offers compared to their counterparts who were PEER, female, and transgender (13). Furthermore, at present there is an underexplored focus of analysis of how gender and gender identity impact the number of faculty job offers a PhD holder may receive. There has been some initial description of the impact (16,17), but it still requires additional work.

Our research is interested in determining factors that lead to success on the faculty job market (2,10,11). This study builds and expands our previous work in the fields of biological sciences to incorporate how PEER and transgender/gender non-conforming (GNC) status impact faculty job offers. We have also included modelling data from our respondents that has characterized what specific factors resulted in job offers from our respondents. Data were collected by self-reported surveys after three job cycles (2019 to 2020, 2020 to 2021, and 2021 to 2022) measuring gender, undergraduate institution type, number of postdoctoral positions, career transition awards, publications, fellowships, number of applications submitted, and the field of the applicants. Each cycle lasted from spring and closed the following year in September.

## 2 Materials and methods

### Ethics

Participation in surveys was voluntary and the respondents could choose to stop responding to the surveys at any time. The three ‘Job Applicant’ surveys were verified by the University of North Dakota Institutional Review Board (IRB) as Exempt according to 4 5CFR46.101(b)(2): Anonymous Surveys No Risk on 08/29/2019. IRB project number: IRB-201908-045.

### Data collection

Individuals from the biological sciences field were included in this analysis. We designed a survey to collect self-reported demographics and academic metrics for assistant professor applicants during the 2019 to 2020, 2020 to 2021, and 2021 to 2022 academic job search cycles. These surveys were open from May to September of each cycle. Respondents were not required to answer all questions. Variables of interest for this analysis included faculty application process outcomes such as interviews, offers and their corresponding institutions; applicant offer responses; and applicant demographics including gender, race, research category, and position. Respondents were also asked to report several productivity metrics including the number of peer-reviewed papers, first-author peer-reviewed papers, *Cell/Nature/Science* (CNS) papers, and first-author CNS papers they published as well as their Google Scholar citation number and h-index. Respondents were also asked about research funding, including post-doctoral fellowship grants.

The survey was distributed on social media platforms including the Future PI Slack group, Twitter, and Facebook as well as by several postdoctoral association mailing lists in North America and Europe. Survey responses that did not meet the minimum completion threshold of 33%, indicated the respondent had previously held a tenure-track position, or did not report submitting any applications were dropped from the analysis. Only respondents who self-categorized their research as biology were included in this analysis.

Aggregated data and survey questions are available in the GitHub repository: https://github.com/Faculty-Job-Market-Collab/Jadavji_Biomed_Frontiers_2024.

### Data categorization and analysis

Where institutions were named, the institution names were cleaned manually and joined with the 2018 Carnegie classification data (https://carnegieclassifications.iu.edu/downloads.php). Using these data, we classified educational institutions based on the National Science Foundation definition for primarily undergraduate institutions (PUIs). PUIs were classified as colleges and universities that awarded 20 or fewer Ph.D./D.Sci. degrees during the previous academic year. An institution with more than 20 Ph.D/D.Sci. degrees awarded during the previous academic year was classified as a research intensive (RI) institution.

Respondents were grouped into three gender-based categories: Man, Woman, and Trans/Gender non-conforming (LGB+/GNC). Respondents were also grouped into two race/ethnicity-based categories: Persons historically excluded due to ethnicity or race (PEER) and non-PEER (13). Respondents were allowed to select as many identities as appropriate from the following list: (i) African-American/Black/African, (ii) Asian-American/Asian, (iii) Caucasian-American/European, (iv) Caucasian-American/North African or Middle Eastern, (v) North American Hispanic/Latinx, (vi) South/Central American, (vii) Caribbean Islander, (viii) North American Indigenous, (ix) Oceania, and (x) Not Listed. Respondents who identified only as Asian-American/Asian, Caucasian-American/European, and/or Caucasian-American/North African or Middle Eastern were considered non-PEER individuals whereas those who selected at least one of the remaining seven race/ethnicity options were considered PEER individuals.

Data were manipulated and visualized using R statistical software (version 4.2.2) and relevant packages. The Pearson’s Chi-squared test with simulated p-values was used to compare respondent demographics and application outcomes. Wherever statistical analyses were used, the tests are reported in the corresponding figure legend. A p-value of less than 0.05 was considered significant. All code used for data analysis and visualization are available in the GitHub repository: https://github.com/Faculty-Job-Market-Collab/Jadavji_Biomed_Frontiers_2024.

### Prediction of faculty application outcomes using machine learning approaches

Here, we predicted faculty application outcomes in the biological sciences participants from all three cycles (2019–2022) based on relevant variables mentioned above, including: gender, age, residence, disability, first-generation undergraduate status, PEER status, number of dependents, position at the time of application, number of postdocs, number of application cycles, number of first-author papers, number of peer-reviewed papers, number of Google Scholar citations, and Google Scholar h-index.

We first standardized the features using the StandardScaler function from the Python scikit-learn library (v 1.2.2), ensuring a mean of 0 and a standard deviation of 1. Then we trained a logistic LASSO (least absolute shrinkage and selection operator (LASSO)) regression model (18) with the above features and binary application outcome labels (i.e., received an offer or not) also using the Python scikit-learn library (v 1.2.2). This model was selected due to potential multicollinearity among independent variables. A 10-fold cross-validation was employed using the KFold method to ensure the robustness of our results. The logistic regression model was configured with the ‘liblinear’ solver, a maximum of 1000 iterations, and a penalty parameter C=1. The average accuracy and its standard deviation across the folds were calculated using the cross_val_score function. For hyperparameter tuning, GridSearchCV was utilized to find the optimal penalty parameter *C* and the maximum number of iterations. The best estimator was identified and validated using the area under the receiver operating characteristic curve (AUROC) as the scoring metric.

To evaluate the model performance, we reported the AUROC, sensitivity, and specificity. Further, we reported the feature importance ranking (based on odds ratio) to understand the predictive power of each variable in the model. Feature importance was visualized using a bar plot of the non-zero coefficients from the logistic regression model. All modeling analyses were conducted in Python (v 3.10.12). All code used for data analysis and visualization are available in the GitHub repository: https://github.com/Faculty-Job-Market-Collab/Jadavji_Biomed_Frontiers_2024.

## Results

We designed a survey for early-career researchers aimed at bringing transparency to the academic job market. Respondents in biological sciences were included in this study. The survey was distributed via Twitter, the Future PI Slack group, and email listservs of multiple postdoctoral associations. The resulting 449 responses (Table 1) were from self-identified early-career researchers in the biological sciences who applied for academic positions in the 2019 to 2020 (n = 231), 2020 to 2021(n = 64), and 2021 to 2022 (n =154) application cycles. This data is also available on our Faculty Job Market Collaboration Dashboard, https://faculty-job-market-collab.org/dashboard/

**Table 1.** Demographics of biological sciences faculty job market survey respondents after the 2019 to 2022 job cycles.

|  |  | n | % |
| --- | --- | --- | --- |
| Total respondents |  | 449 |  |
| Gender | LGB+/GNC | 61 | 13.12 |
|  | Man | 150 | 32.26 |
|  | No response | 1 | 0.22 |
|  | Woman | 253 | 54.41 |
| Age | 26 to 30 years old | 36 | 7.74 |
|  | 31 to 35 years old | 251 | 53.98 |
|  | 36 to 40 years old | 146 | 31.4 |
|  | 41+ years old | 32 | 6.88 |
| Dependents | No | 305 | 65.59 |
|  | Yes, one child | 83 | 17.85 |
|  | Yes, multiple children/adult(s) | 77 | 16.56 |
| Disability status | No | 400 | 86.02 |
|  | No response | 12 | 2.58 |
|  | Yes | 53 | 11.4 |
| First Generation |  |  |  |
| PhD Student | No | 93 | 20 |
|  | Unsure | 3 | 0.65 |
|  | Yes | 369 | 73.98 |
| Undergraduate | No | 344 | 73.98 |
|  | Unsure | 4 | 0.86 |
|  | Yes | 117 | 25.16 |
| Position at time of survey | Non-tenure track faculty | 59 | 12.69 |
|  | PhD candidate (ABD) | 3 | 0.65 |
|  | Postdoc | 318 | 68.39 |
|  | Tenure track Assistant Professor | 46 | 9.89 |
|  | Not applicable | 39 | 8.39 |
| Residence | Canada | 29 | 6.24 |
|  | Other | 24 | 5.16 |
|  | USA | 412 | 88.6 |
| Legal status | Citizen/resident | 370 | 79.57 |
|  | Not applicable | 1 | 0.22 |
|  | Other | 10 | 2.15 |
|  | Visa | 84 | 18.06 |

Abbreviations: ABD, all but dissertation; GNC, gender non-conforming (e.g. transgender, gender fluid); LGB+, lesbian, gay, bisexual, plus.

### Demographics of respondents

Most respondents were women (54.41%, Table 1) and between the ages of 31 to 35 (53.98%). A small (13.12%) percentage of respondents were LGB+/GNC. Furthermore, a small portion (34.41%) of respondents had dependents, and most were first generation graduate students (79.35%). A subset of respondents indicated a disability status (13.98%). Applicants were mostly postdocs (68.39%) and from the US (88.6%). We did collect responses from Canadian residents (6.24%) as well as those with other countries of residence (5.16%).

### Respondent application process

Through our survey we collected several academic metrics that we have previously examined in order to generate a longitudinal data set, which has facilitated the creation of our data dashboard (2,10,11). Respondents reported a wide range in the number of submitted applications from minimum of 1 to a maximum of 96 (median: 15; Figure 1). The median number of faculty job offers was 1 with a range of 0 to 8. Applicants had a median number of 6 first-author papers with a median of 11.5 peer reviewed papers and 1 corresponding author paper. The median number of citations for candidates was 355 with a range of 4 to 6585. The most common number of citations (mode) was 935. The median h-index (as obtained through Google Scholar) of respondents was 8 with a range of 0 to 23. In figure 1 (second column) we also include metrics of respondents that obtained at least one job off.

**Figure 1.**
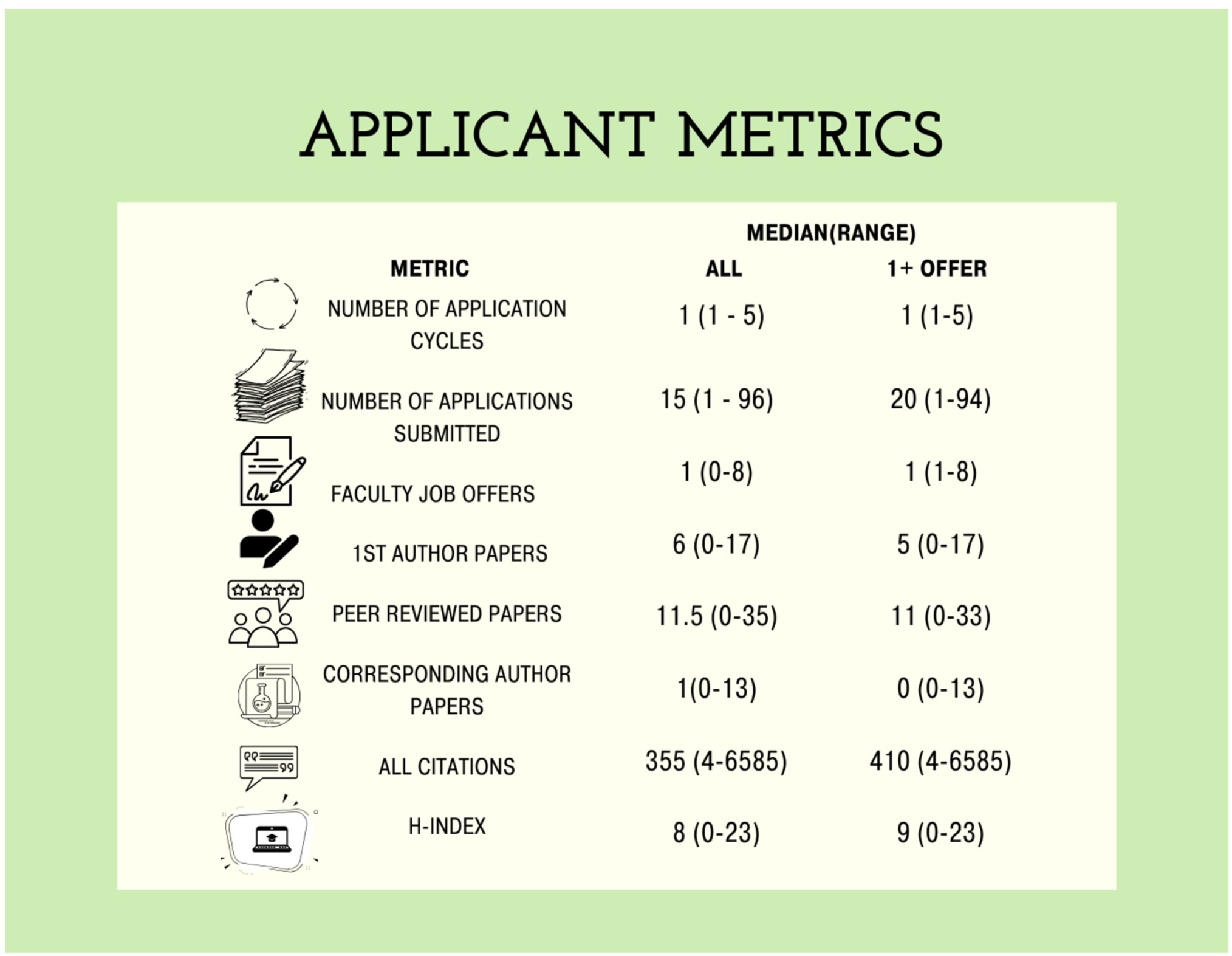
Applicant median and range values for metrics of research productivity in biological sciences respondent pool. Values include data from all respondents and individuals that received at least one job offer.

In terms of funding, most respondents were not listed as a principal investigator (PI; 76.34%) or Co-PI (82.15%) on a research project grant (Figure 2). A small portion (4%) of respondents that received job offers that help research grants as a PI.

**Figure 2.**
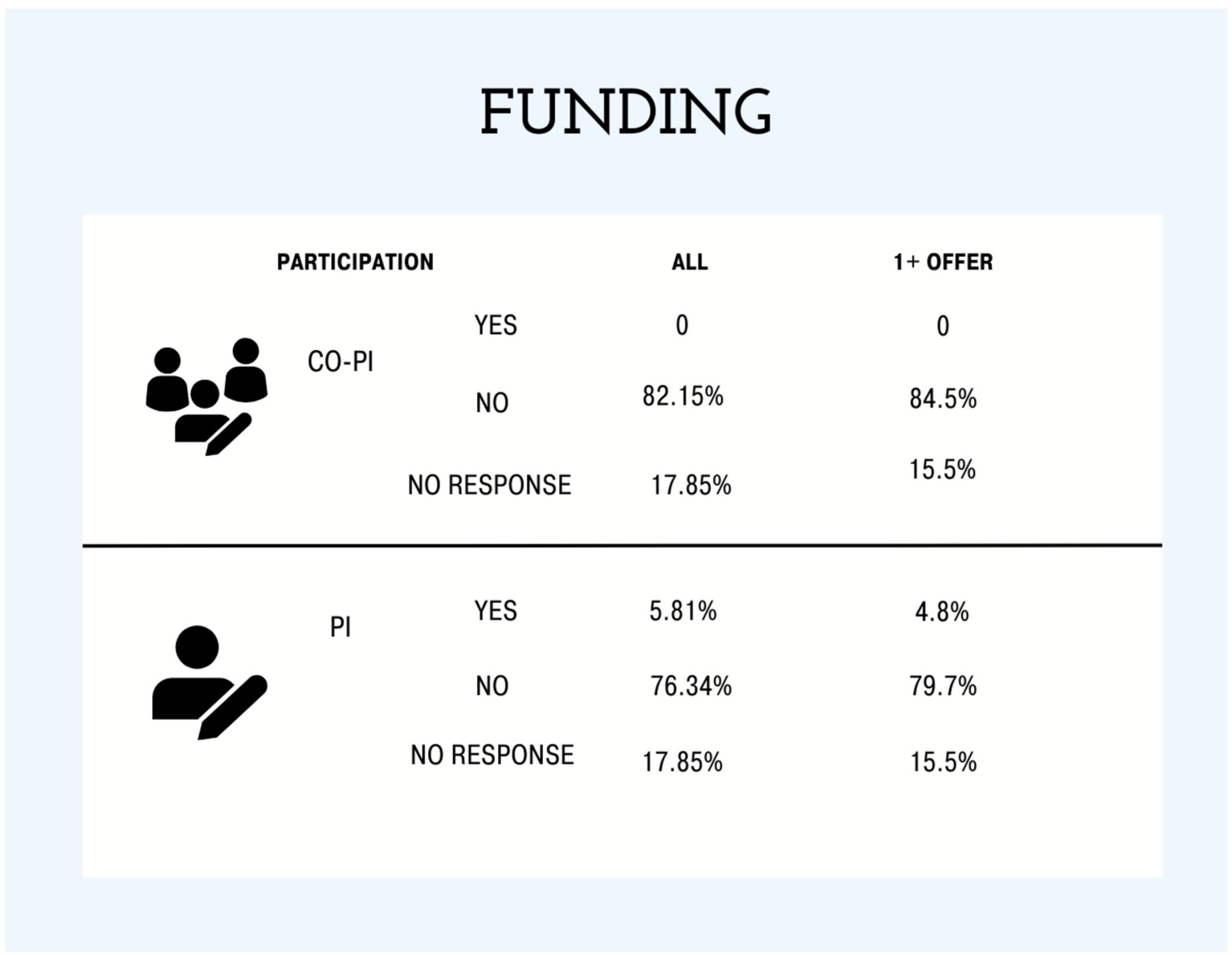
The percentage of research funding project obtained as principal investigators (PI) or co-principal investigator (co-PI) of all respondents and individuals that received one or more faculty job market offer(s).

### Impact of PEER status on the academic job market

Most respondents to our survey did not select ethnicities that met our definition of a PEER status (Figure 3A) (13). We did investigate whether there were differences between PEER and non-PEER groups in terms of some scholarly metrics and success on the faculty job market. There was no difference in the total number of applications submitted for faculty jobs between PEER and non-PEER groups (Figure 3B). However, both PEER and non-PEER status PhD holders applied to more RI institutions compared to PUIs (Figure 3C, p = 0.000099). There was no difference in the number of *CNS* first author papers (Figure 3D), total number of *CNS* papers published (Figure 3E), or number of onsite interviews (Figure 3F) between PEER and non-PEER groups. Overall, we did not observe differences in the number of faculty offers between PEER and non-PEER groups (Figure 3G). Additionally, we report that in our survey respondent having PEER status and being a first-generation undergraduate student was positively correlated (r = 0.219, p = 0.0001). This positive correlation was present for PhD first-generation students and having PEER status, but was not significant (r = 0.006, p = 0.207).

**Figure 3.**
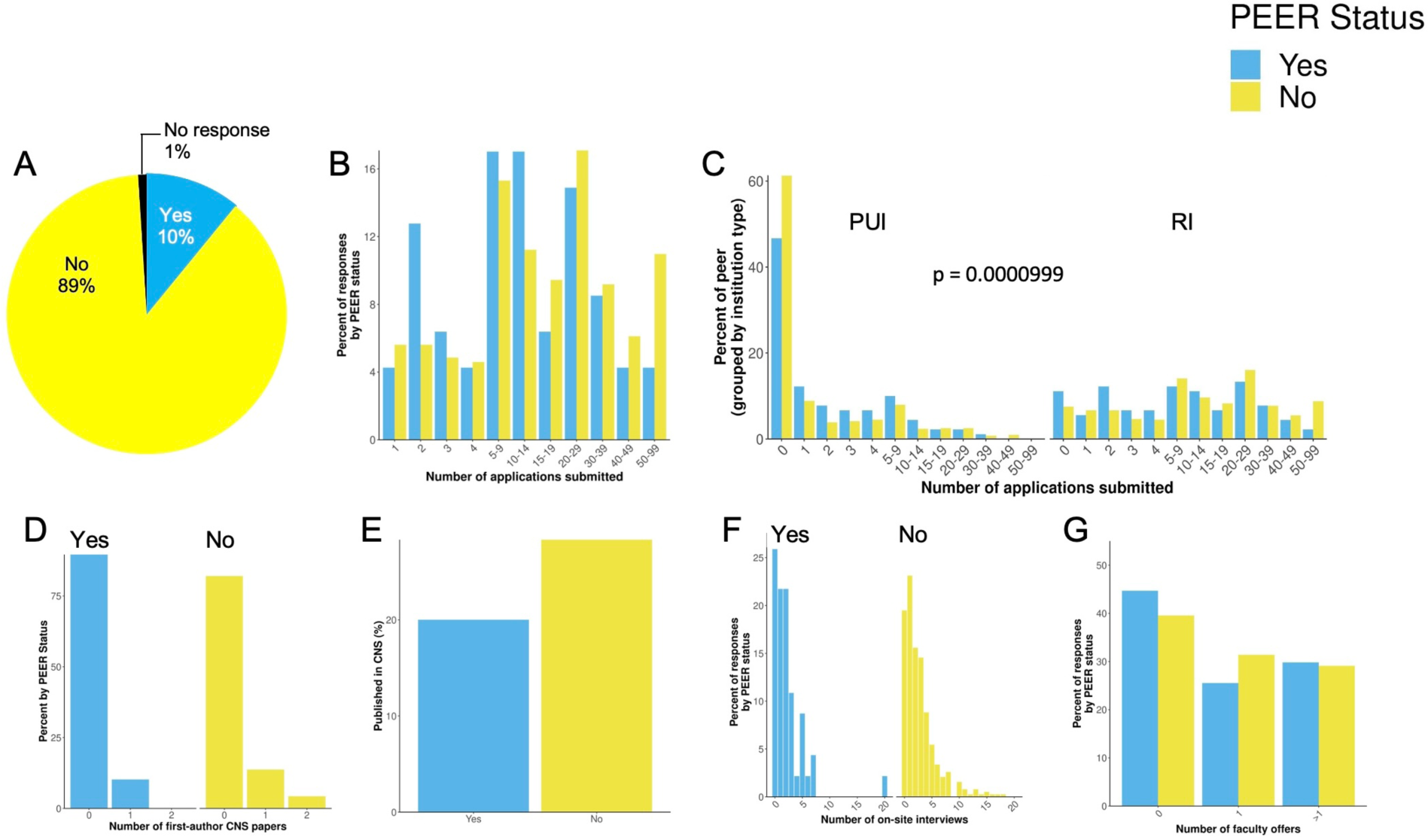
Respondent data organized for persons excluded because of their ethnicity or race (PEER) status and non-PEER status. A, percentage of respondents that self-selected ethnicities that met our definition for PEER status. B, the number of applications submitted. C, number of applications submitted by institution type (primarily undergraduate, PUI and research intensive, RI). D, number of first author *Cell*, *Science*, or *Nature* (CNS) publications. E, total number of CNS publications. F, the number of onsite interviews and G, the number of faculty offers. Respondents with PEER and non-PEER status submitted more applications to research intensive (RI) institutions compared to primarily undergraduate institutions (PUI; p = 0.0000999).

### Impact of gender on the academic job market

As mentioned above, many of our survey respondents identified as women (54.41%, Table 1). We did not see a difference in the number of applications submitted (Figure 4A) between men, women, and LGB+/GNC respondents. Most applicants, regardless of gender, reported applying for positions at RI institutions compared to PUIs (p = 0.0000499, Figure 4B). There was no difference in the percentage of publications in *CNS* (Figure 4C), but women and LGB+/GNC had fewer first-author publications in *CNS* (Figure 4D, p = 0.01129). There was no difference between genders in the number of onsite interviews (Figure 4E). Overall, our data showed that women had more job offers (Figure 4F, p = 0.0229).

**Figure 4.**
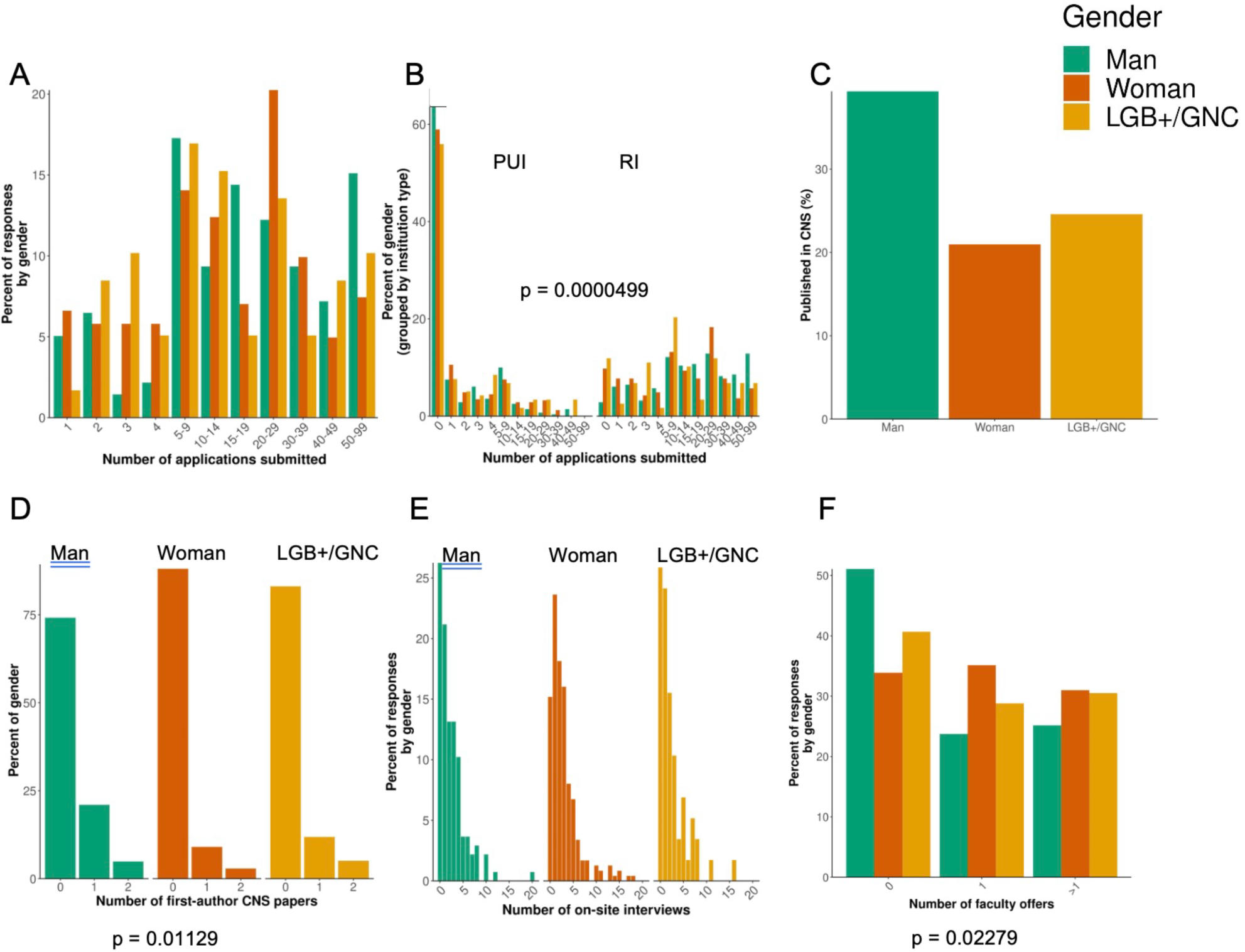
Respondent data organized by gender. A, the number of applications submitted. B, number of applications submitted by institution type (primarily undergraduate, PUI and research intensive, RI). C, number of first author *Cell*, *Science*, or *Nature* (CNS) publications. D, total number of CNS publications. E, the number of onsite interviews and F, the number of faculty offers. All genders submitted more applications to RI institutions compared to PUI (p = 0.0000499). Respondents that were women, LGB+ (lesbian, gay, bisexual, plus) or gender non-conforming (GNC). LGB+/GNC had less first author CNS papers. Furthermore, respondents that were women had more job offers compared to men or LGB+/GNC.

### Factors that predict success on the academic job market

We implemented the LASSO logistic regression model using data from all three cycles (2019 to 2022) (n = 449; 287 or 63.92% received offers). The model achieved an area under the receiver operating characteristic curve (AUROC) of 0.68 (95% C.I.: 0.53 – 0.83), with a sensitivity of 0.84 at a specificity of 0.36, suggesting that the model is better at identifying candidates who would successfully receive a job offer than excluding those who would not. The feature importance descends in the following order (largest effect first): number of applications (positive), senior position at the time of application (positive), number of postdocs (negative), gender (positive, woman = 1, men = 0), whether is first generation undergraduate (negative), age (negative), and residence (negative, reside out of U.S. or C.A. = 1) (Figure 5). In summary, job candidates in biological sciences who submitted more applications in the most recent cycle, had a more senior academic position, and/or were women or LGB+/GNC had higher odds of receiving a faculty offer; in contrast, job candidates who conducted more rounds of postdocs, were first generation undergraduate, older in age, and reside outside of U.S. or C.A. had lower odds of receiving a faculty offer.

**Figure 5.**
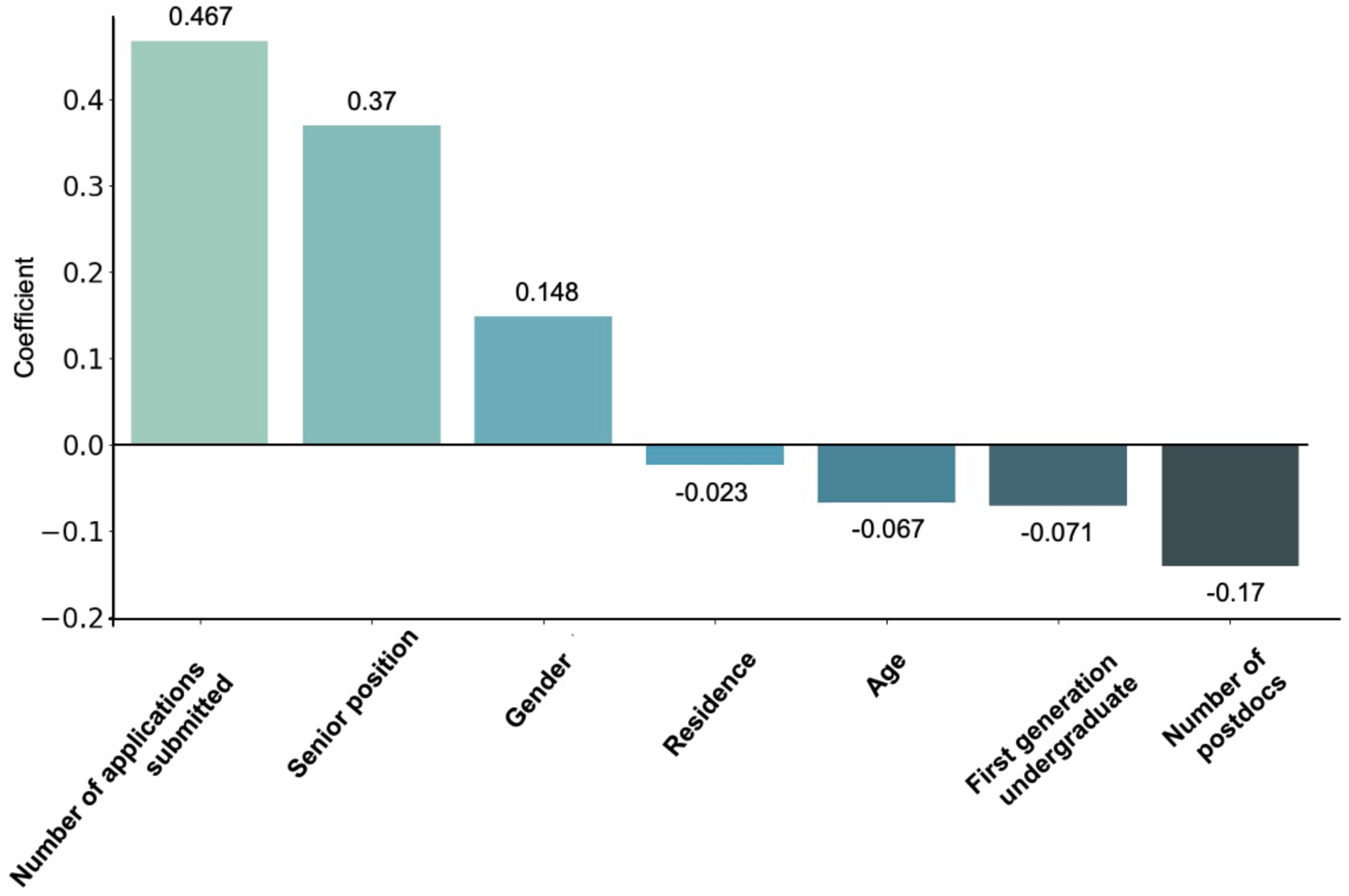
Regression model analysis showing the relationship between factors and receiving faculty job offers in the biological sciences over three cycles (2019 to 2022). We report a positive relationship following order (largest effect first): number of applications submitted, senior positions, and gender (identifying as a women). Negative relationships are reported for residence, age, first generation undergraduate status, and the number of postdocs.

## 3 Discussion

In the biological sciences the faculty job market is elusive, and most PhD holders have an incomplete understanding of what is required to be successful nor when they should go on the market. To shed some light on factors that make individuals competitive for faculty positions in the biomedical field, we started a national collaboration in 2018 (11). This current study is a continuation of our initial work. In addition, we have also expanded our research to include disability and PEER status, as well as LGB+/GNC individuals. We surveyed individuals who were on the faculty job market for a biological sciences position after three cycles spanning 2019 to 2022. Our present study has shown that more job applications submitted results in more offers. Additionally, we have also shown that there is not a significant impact of PEER status on obtaining a faculty job offer in the biological sciences. We also show that women respondents obtained more job offers, and men respondents had more CNS first-author papers. Furthermore, our results show that increasing the number of postdoc positions did not result in more job offers.

Our results from this study are consistent with data from our previous work in 2018, which showed that the number of job applications submitted is positively correlated with the number of job offers received (11). Most faculty job market applicants are encouraged to apply broadly (19). Our data for now four (2018 to 2019, 2019 to 2022) job cycles has shown that more applications submitted is positively correlated with a job offer (11). Furthermore, faculty hiring is not standardized, and search committees sometimes have vague (20) research classification or change their direction in research areas based on applications they received (17,21). PhD holders that are on the faculty job market may consider broadly applying to jobs to increase the number of the applications they submit and consider that sometimes job ads are left vague on purpose.

Our survey is voluntary to begin, and all questions are optional. Interestingly, we report that women had more job offers than men. In the present study, we had more women respondents (54.4%) than men. Other survey respondent studies have shown that women are more likely to complete surveys online (22). Recruiting all three gender categories for future surveys will help remove this bias in the data.

Our modeling data show that the number of postdocs was found to correlate with lower faculty job application success. Of the 449 respondents, 68.39% of them were postdocs at the time of survey completion, while less than 10% were hired as tenure-track assistant professors. These results were unexpected because of the assumption that a longer or greater number of postdocs would increase the competitiveness of the application (7). It is possible that postdocs who were unsuccessful in their academic job search were more likely to participate in the survey. Other negative coefficients were first generation undergraduates, age, and gender. Further research into the PhD job market should account for additional extrinsic factors to get a more complete understanding of the evolving job market and should include a greater number of faculty respondents to eliminate any bias in the data.

In the present study, we expanded our survey to include persons excluded because of their ethnicity or race (PEER) status. When comparing the number of faculty applications PEER and non-PEER status PhD’s, both groups submitted a comparable number, with PEER and non-PEER counterparts both submitting more applications to RI institutions. In terms of academic and career merits, non-PEERs had a greater percentage of first authorship CNS papers, a lower number of on-site interviews, and a higher number of faculty offers, it is important to note that these were trends in the data and not mathematically different. These findings may suggest that PEER status can affect the success of a candidate’s faculty job market application, even when other academic metrics exceed those of non-PEER applicants. Additionally, we report that most PEER status respondents were first-generation college undergraduate students, which is interesting, this positive correlation was not as strong for PhD holders. Our work has generated some interesting data in relation to PEER status, but also suggests that the experiences of PEER faculty applicants should be studied in more depth.

We have continued to provide more insight into what factors play an important role in landing a faculty job in the biological sciences. The data from this survey is available online through our dashboard (https://faculty-job-market-collab.org/dashboard). We strongly believe that further data collection is vital for future trainees, advisors, and administrators to help facilitate career development and decision making regarding whether to go on the job market. However, it is likely that numbers and metrics alone do not fully capture the dynamics of the process. Each PhD holder that participated in the faculty job market has a unique trajectory, perceptions, and experiences on the market that is not captured by our quantitative data collection, therefore compiling these experiences into a qualitative analysis would further enrich our understanding and provide relevant insight to efforts to broaden participation in academic biological and biomedical science.

## 5 Author Contributions

Conceptualization: A.J.K, A.K.H., C.T.S., A.H., N.M.J., Data Curation: A.K.H., Formal Analysis: A.K.H., Y.C., Funding Acquisition: A.J.K, Y.C., A.K.H., C.T.S., A.H., N.M.J., Investigation: A.J.K, Y.C., A.K.H., C.T.S., A.H., N.M.J., Methodology: A.K.H., Y.C., Software: A.K.H., Y.C., Supervision: A.H., N.M.J., Visualization: A.J.K., A.K.H., Y.C., Writing – original draft: B.F., A.J.K, Y.C., A.K.H., C.T.S., A.H., N.M.J., Writing – review and editing: B.F., A.J.K, Y.C., A.K.H., J.N., C.T.S., A.H., N.M.J.

## 7 Acknowledgments

We would like to thank Google Cloud for providing cloud computing resources through their free tier, which supported the predictive modeling in this study.

## 8 Supplementary Material

None

## 9 Data Availability Statement

Faculty Job Market Collaboration Data Dashboard, https://faculty-job-market-collab.org/dashboard/

Faculty Job Market Collaboration generated and anonymized dataset, https://github.com/Faculty-Job-Market-Collab/Jadavji_Biomed_Frontiers_2024

